# Engineered lymphatic stroma model applications in central nervous system leukemia

**DOI:** 10.1101/2025.08.13.670203

**Authors:** Jennifer H Hammel, Peter M Gordon, L. Monet Roberts

## Abstract

Central nervous system (CNS) involvement frequently occurs in acute lymphoblastic leukemia (ALL), but many questions remain about how leukemia cells access, persist in, and exploit the CNS. The CNS is protected by the meninges, which are fluid filled membranes that surround the brain. Within the meninges, meningeal lymphatic vessels drain cerebrospinal fluid to cervical lymph nodes for immunosurveillance, a potential pathway for leukemia cell migration into or egress from the CNS. Here, we utilize tissue engineered models of the meningeal lymphatics and lymph node stroma to probe how leukemia cells interact with these key tissues. We first demonstrate that standard-of-care chemotherapeutics can damage meningeal lymphatic barriers. Next, we showed that soluble factors from the meningeal lymphatics can support leukemia cell growth while soluble factors from the lymph node model under flow can promote leukemia cell migration. Finally, we show that leukemia cells migrate through the lymph node model under both static and flow conditions. Overall, we have demonstrated the feasibility of using engineered lymphatic models to study leukemia cell behavior in the CNS with the goal of expanding the available experimental platforms for understanding CNS metastasis and relapse.

**Insight Box:** Leukemia commonly infiltrates the central nervous system (CNS), requiring intensive CNS-directed therapies that are often ineffective and cause both acute and long-term toxicities, especially in pediatric patients. The meningeal lymphatics and the deep cervical lymph nodes constitute a pivotal axis in CNS immunity, facilitating drainage of fluid and waste and enabling peripheral immune surveillance in response to CNS-derived signals. Here, we employ in vitro models of the meningeal lymphatics and lymph node stroma to demonstrate their crosstalk in influencing leukemia cell growth and migration. These engineered platforms serve as valuable tools for uncovering mechanistic insights into the meningeal-lymph node axis in the context of CNS-leukemia relapse.

## Introduction

Acute lymphoblastic leukemia (ALL) is a hematologic malignancy in which immature lymphoid precursor cells fail to differentiate into functional white blood cells. These leukemia cells originate in the bone marrow but can disseminate through the bloodstream and infiltrate distant organs. The central nervous system (CNS) is a particularly critical site of leukemia involvement and a common site of relapse. Though CNS-targeted therapy has decreased overall rates of relapses significantly, the CNS is still involved in 30-40% of all relapses (1).

Current therapeutic strategies for CNS leukemia include cranial irradiation and chemotherapy (2). Cranial irradiation is utilized sparingly, due to the significant side effects on the developing brain of pediatric patients. Chemotherapeutic agents, methotrexate and cytarabine, are typically administered intrathecally, or into the cerebrospinal fluid, to target the meninges and CNS. In addition, both drugs can cross the blood-brain barrier when given systemically at high doses (3,4). Though these agents are effective, studies have shown long-term adverse neurocognitive effects after treatment (5–7). These CNS-directed therapies result in chemotherapy treatment directly applied to the meninges. Notably, recent studies have implicated the meninges in CNS relapse by promoting leukemia cell quiescence and conferring resistance to chemotherapy (8). Within the meninges, lymphatic vessels provide nutrients, facilitate waste clearance, support fluid homeostasis, and drain cerebrospinal fluid to the deep cervical lymph nodes. The meningeal lymphatics have been shown to contribute to the efficacy of radiotherapy in the treatment of glioblastoma (9), and we have previously demonstrated chemotherapy-mediated disruption to the meningeal lymphatics (10). However, it is unknown how the meninges and meningeal lymphatics may be involved in the efficacy and off-target effects of CNS-directed chemotherapy.

Additionally, the route that leukemia cells utilize to traffic in and out of the meninges is unclear (11). Drainage into the cervical lymph nodes from the meningeal lymphatics subsequently facilitates immune cell trafficking from the periphery, particularly in response to inflammatory stimuli (9,12,13). Moreover, peripheral immunosurveillance has recently been shown to exploit the meningeal lymphatic- lymph node axis in response to radiation therapy (9). Given these important mechanisms, it remains unknown whether this same route is also engaged in leukemia cell trafficking, co-opting tumor-specific immune responses during disease progression (11). Further, lymphatic endothelial cells (LECs), one of the major components of the lymphatic vasculature, secrete a number of cytokines that act as a chemoattractant for immune cells and tumor cells (14–16), potentially attracting leukemia cells to intravasate into the vasculature as an unintended effect. Once in the circulation, these cells also encounter various fluid flows as they travel within the vasculature with leukemia cells migrating in the direction of shear stress (17). Together, LEC-derived chemokines and physical extracellular cues are important in potentially mediating leukemia cells along drainage pathways towards the lymph node.

The lymph node is a complex and dynamic microenvironment with constant immune cell recirculation and interstitial fluid flow (IFF) delivering pathogens for immunosurveillance (18,19). In leukemia, once tumor cells reach the lymph node, they can be disseminated via the lymphatic system and cause relapse in distant organs in the body. In support of this, the lymph node microenvironment has been implicated in regulating ALL cell proliferation (20), though more in-depth studies of the lymph node have primarily focused on chronic lymphocytic leukemia (21,22),. Additionally, IFF is greatly enhanced during immune response and subsequent inflammation, resulting in immune cell retention. Yet, the impact of IFF on leukemia cells in the lymph node has never been examined. Thus, open questions remain about interactions within the lymph nodes in ALL.

Creating tools to study leukemia cell behavior in the brain to cervical lymph node pathway will provide a platform to develop targeted therapeutics for CNS leukemia. Thus, we sought to apply our established lymphatic stroma models (10,23) toward varying applications in understanding CNS leukemia. Toward that goal, herein we first examined cellular responses to CNS-directed chemotherapy through our simple in vitro model of the meningeal lymphatics (10). We then looked at how the naive and chemotherapy-treated meningeal lymphatic microenvironments can alter leukemia cell growth. To represent downstream interactions within the lymph node, we also utilized our model of the lymph node stroma (23) to assess the intersection of IFF and chemoattraction in leukemia cell egress from the meningeal lymphatics. Collectively, our findings demonstrate that the lymphatics involved in CNS immunosurveillance are an essential consideration in studying CNS leukemia and that our engineered models of the lymphatic stroma are a useful platform toward understanding leukemia growth and migration.

## Methods

### Cell culture

Human lymph node lymphatic endothelial cells (LECs) (Sciencell) were cultured on fibronectin coated flasks in VascuLife® VEGF-Mv media (Lifeline Cell Technology) and human meningeal cells (MCs) were cultured in poly-L-lysine-coated flasks in MenCM (Sciencell) as previously described (10). Jurkat leukemia cells were cultured in RPMI with 10% characterized fetal bovine serum (Cytiva Hyclone).

### Engineered meningeal lymphatic tissue culture model and treatment

LECs and MCs were incorporated into the meningeal lymphatic (ML) model as previously described (10). Briefly, 50,000 LECs were seeded on the underside of an inverted 12-mm 8-micron pore size tissue culture insert (Corning) and re-inverted after 2 hours. After 2 days, 75,000 MCs were added into the tissue culture insert. The model was cultured for an additional 24 hours prior to the initiation of experiments. For chemotherapy treatment studies, ML models were treated with either 500 nM methotrexate (MTX), cytarabine (Ara-C), or vehicle (sodium hydroxide) for 24 hours. MTX was initially dissolved to 1 mg/mL in 0.1M sodium hydroxide and diluted to 500 nM in VascuLife® VEGF-Mv media (Lifeline Cell Technology). The vehicle was adjusted to account for the small amount of 0.1M sodium hydroxide (0.5 µL per mL of media). Ara-C was dissolved in water. Media from treated ML models was flash frozen and stored at -80º for conditioned media assays.

### Immunofluorescence and imaging of chemotherapeutic-treated meningeal lymphatic models

ML models were fixed with 4% paraformaldehyde for 15 minutes at room temperature (RT). The membranes of each insert were cut out and mounted on positively charged slides. Membranes were incubated with 3% donkey serum to block nonspecific binding followed by 4 µg/mL sheep anti-human CD31 (R&D) for 2 hours at RT. Models were rinsed with PBS three times, followed by a 1 hour incubation at RT with 5 µg/mL donkey anti-sheep AlexaFluor 647 in blocking solution. Finally, membranes were stained with 1:400 488 Phalloidin for F-actin (Invitrogen) and 1:5000 DAPI (Invitrogen) at RT for 45 minutes. Slides were mounted with Fluoromount, sealed with nail polish and stored at 4°C until imaging on a Zeiss Axio Observer.

Three regions of interest at 20X were imaged from each membrane with three technical replicates of membranes for a total of 9 images per condition. LEC coverage was quantified via % area CD31 signal in Fiji (24). LECs were traced and area and aspect ratio were recorded from FIJI. Cells with disrupted junctions were counted by hand and reported as a percentage of total cells.

### Jurkat growth curve

Jurkats were plated at 200,000/mL in conditioned ML model media, 500 nM MTX in Vasculife, fresh Vasculife, or fresh RPMI in low attachment 96-well plates. Every 12 hours, a 4 uL sample was taken from each well and counted via a Nexcelom Bioscience Cellometer Automated Cell Counter to track cell count over time.

### Leukemia cell egress from the meningeal lymphatic model

For conditioned media studies, supernatant was collected from lymph node stroma models under static and flow conditions (25) and kept at 4°C until use. Conditioned media was placed in the well underneath the ML model insert and fresh Vasculife was placed within the insert. Jurkat cells labeled with CellTracker Orange (Invitrogen) were added to the upper compartment of the ML model at 50,000 cells per tissue culture insert. After 24 hours, media from underneath the tissue culture insert was collected and Jurkat cells were counted manually.

ML models for imaging were fixed in 4% PFA for 15 minutes. The tissue culture insert membranes were removed via a scalpel and placed on a positively charged slide. ML models were incubated in blocking solution (donkey serum in 0.01% Triton X-100 in PBS) for 1 hour at RT. ML models were then incubated with 1 µg/mL mouse anti-human alpha smooth muscle actin antibody [1A4] (abcam) and 4 µg/mL sheep anti-human CD31 (R&D) in blocking solution for 2 hours at RT. Models were rinsed with PBS three times and then incubated with 2 µg/mL donkey anti-mouse AlexaFluor 555 and 5 µg/mL donkey anti-sheep AlexaFluor 647 in blocking solution for 1 hour at RT. Models were rinsed with PBS three more times and then incubated with DAPI (1:5000) for 15 minutes at RT. Slides were mounted and stored as described above.

### Jurkat invasion assay

Jurkat leukemia cells were encapsulated at 1 million/mL in methacrylated hyaluronic acid (PhotoHA)-collagen in tissue culture inserts. Hydrogel formulation consisted of 0.4% PhotoHA (Advanced Biomatrix) and 0.2% collagen (Corning) as previously described (25). For static conditions, 700 µL of media was placed underneath the tissue culture insert and 100 µL was placed atop the hydrogel for static conditions. For 0.8 µm/s flow conditions, 100 µL was placed underneath the tissue culture insert and 700 µL was placed atop the gel. After 24 hours, hydrogels were removed from the tissue culture insert. The membranes were fixed with 4% formaldehyde in 1X PBS for 15 minutes. Then, membranes were stained with DAPI and three regions of interest were imaged on an EVOS at 10X. Invaded cells were counted and converted to % invasion.

### Leukemia cell egress from the lymph node stroma model

Lymph node stroma models were established as previously described (23). Briefly, LECs were seeded on the underside of a tissue culture insert as described above. Fibroblastic reticular cells (FRCs) are encapsulated in a methacrylated hyaluronic acid (PhotoHA)-collagen hydrogel precursor prior to photo- and thermally crosslinking above the LEC monolayer at 50,000 per insert with subsequent experiments being performed following 24 hours of co-culture.

Jurkat cells were labeled with CellTracker Orange (Invitrogen) and added to the upper compartment of the lymph node stroma model at 50,000 cells per tissue culture insert under static or flow conditions. After 24 hours, media from underneath the tissue culture insert was collected and Jurkat cells were counted manually. Lymph node stroma models were fixed in 4% formaldehyde in 1X PBS for 45 minutes at RT. Blocking with donkey serum in 0.01% Triton X-100 was performed for 6 hours at RT on a rocker. Lymph node stroma models were then incubated with 1 µg/mL mouse anti-human alpha smooth muscle actin antibody [1A4] (Abcam) in blocking solution overnight at 4°C on a rocker. Lymph node stroma models were rinsed with PBS three times for 20 minutes on a rocker at RT. Next, gels were incubated with 2 µg/mL donkey anti-mouse AlexaFluor 555 in blocking solution overnight at 4°C on a rocker, followed by a rinse as previously described. Finally, gels are incubated with DAPI (1:5000) for 45 minutes at RT. Gels are imaged on a Nikon Spinning Disk by placing the tissue culture insert onto a coverslip. Z-stacks were taken at 10X with a 10 micron step size.

### Statistical analysis

Statistical analysis was performed in GraphPad Prism 10.2.2 (GraphPad Software). To determine significance, one-way ANOVA was performed with a p value of <0.05. If significant, post hoc Tukey’s t-tests for multiple comparisons were performed between groups. All graphs represent the mean with each biological replicate displayed.

## Results

### Methotrexate treatment disrupts LECs, but not MCs, in a model of the meningeal lymphatics

Leveraging insights from our previous studies in taxane chemotherapy (10), we first sought to examine whether standard-of-care chemotherapeutics for treating CNS leukemia would cause disruption to the meningeal lymphatics. To answer this question, we utilized our human engineered ML model (10) consisting of LECs and MCs on either side of a tissue culture insert as previously demonstrated (**Figure 1A**). This model represents the lumen of a meningeal lymphatic vessel with the upper compartment representing the stromal component of the meninges and the lower compartment representing the inside of the lymphatic vessel endothelium. Methotrexate (MTX) primarily inhibits leukemia cell proliferation by targeting dihydrofolate reductase, a key enzyme in one-carbon metabolism. In addition to its antiproliferative effects, MTX also exerts notable anti-inflammatory properties.(26). Cytarabine (Ara-C) induces leukemia cell death by impairing DNA synthesis and repair (27). In response to MTX and Ara-C treatment, we found that MCs were not disrupted by either chemotherapeutic agent (**Figure 1B**). However, LEC barrier integrity was impacted. We saw significantly decreased LEC coverage (**Figure 1C**), and significantly increased junction disruption (**Figure 1D**) by MTX but not Ara-C. Next, we sought to further quantify morphological changes on a cell-to-cell level. However, average LEC area (**Figure 1E**) and aspect ratio (**Figure 1F**) were unchanged by chemotherapy treatment.

**Figure 1.**
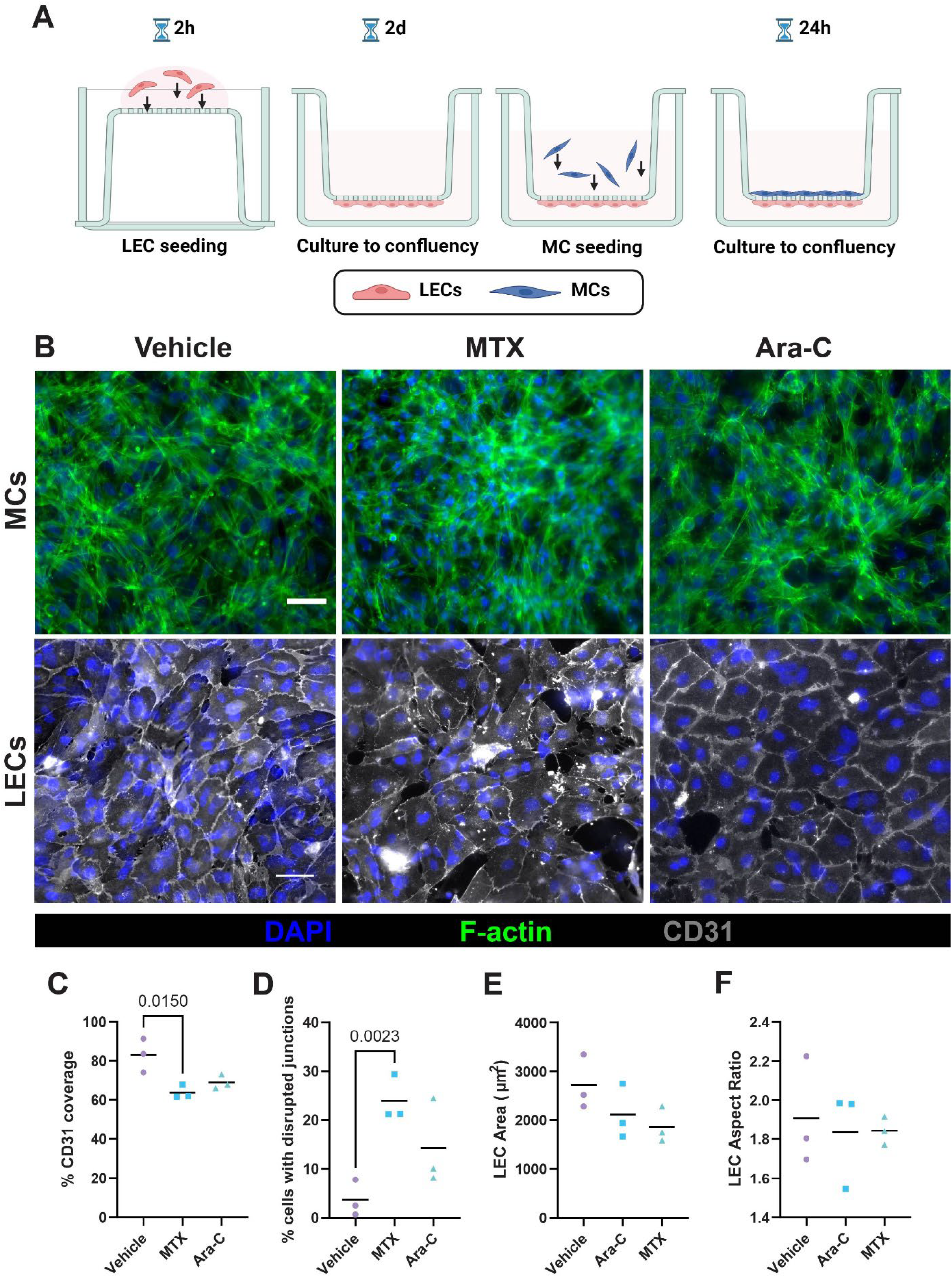
Methotrexate treatment is disruptive to lymphatic endothelial cells within the engineered meningeal lymphatic model. **(A)** Schematic representing the meningeal lymphatic in vitro model with lymphatic endothelial cells (LECs) seeded on the underside of a tissue culture insert and grown to confluency. After two days, meningeal cells (MCs) were added inside the insert and grown to confluency. The model was then treated with either 500 nM methotrexate (MTX), cytarabine (Ara-C), or vehicle. **(B)** Representative images of MCs (top) and LEC (bottom) monolayers with MCs visualized with F-actin stain in green, LECs stained for CD31 in gray and nuclei stained with DAPI in blue. Scale bars are 50 µm. **(C-F)** LEC coverage, disrupted junctions, average area, and average aspect ratio are reported. Data shown represents the mean with each biological replicate shown (n=3). Significance was determined via one-way ANOVA between the vehicle and chemotherapeutic treatments with post hoc Tukey’s multiple comparison test.

### Meningeal lymphatic model conditioned media encourages Jurkat growth

MCs have previously been shown to confer chemoresistance via reduced leukemia cell growth, suggesting a mechanism that promotes relapse in the meninges (8). Thus, we aimed to link chemotherapy-mediated changes observed with cells within the meningeal lymphatics with altered leukemia cell growth. To address this, we collected conditioned media from the chemotherapy-treated ML models and quantified the growth of Jurkat leukemia cells (**Figure 2A**). As expected, MTX conditions demonstrated a trend of higher doubling times, though not significant (**Figure 2B**). Interestingly, Jurkats in the ML conditioned media had the lowest trend in doubling time. When examining the endpoint at 48 hours, the ML model condition had significantly more Jurkats than the RPMI control (**Figure 2C**). Growth curves demonstrate that the ML condition trended higher than MTX conditions and Vasculife controls (**Figure 2D**). These results suggest that the meningeal lymphatics secrete important soluble factors that support Jurkat growth and survival. However, soluble factors through conditioned media do not recapitulate the cell-cell contact between meningeal cells and leukemia cells that is known to impact chemoresistance (8).

**Figure 2.**
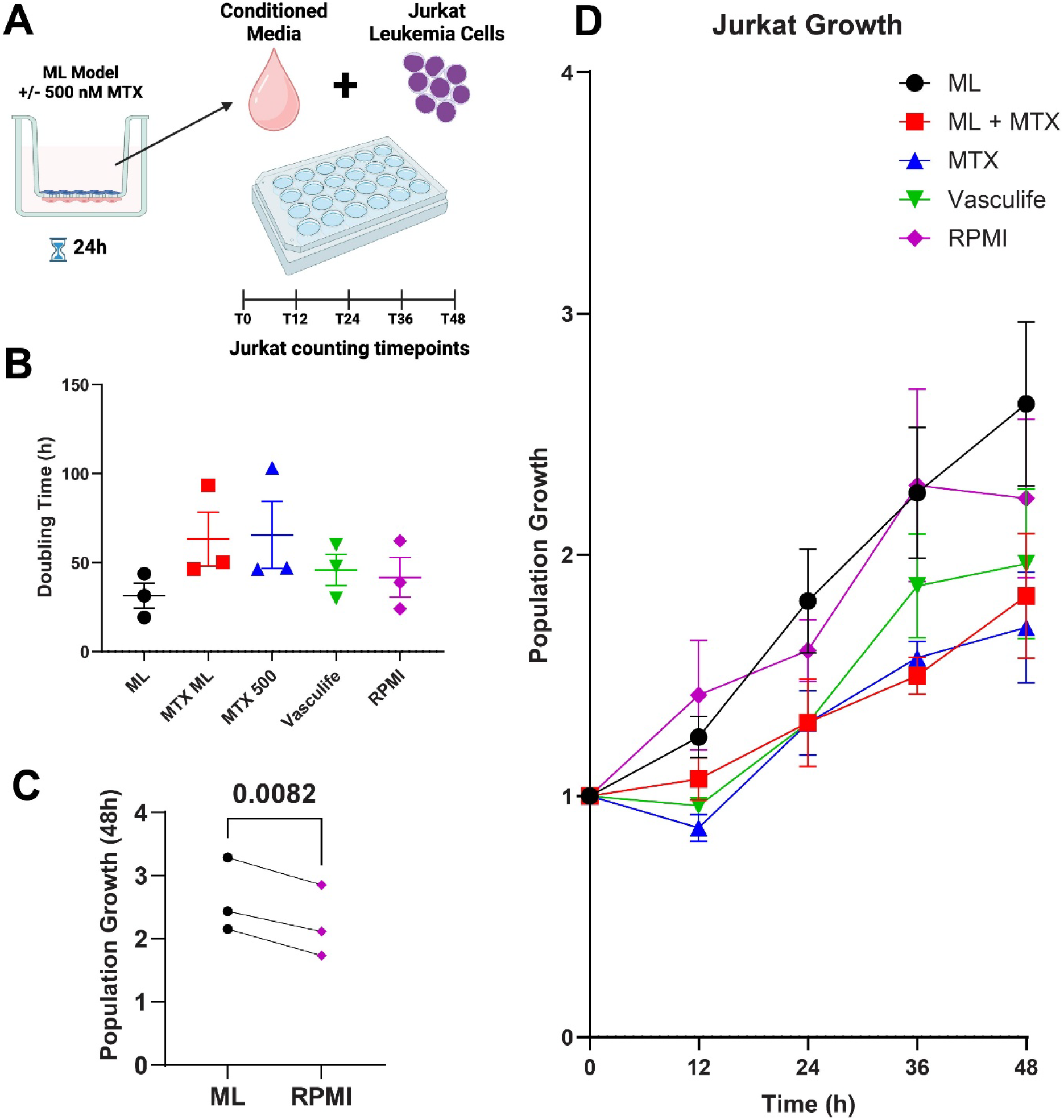
Jurkat growth trends are altered by ML model conditioned media. **(A)** ML models were treated with 500 nM methotrexate for 24 hours before media was collected and stored. ML represents ML model conditioned media. MTX 500 represents fresh Vasculife with 500 nM MTX as a control. Jurkats were grown in conditioned media for 48 hours and counted every 12 hours. **(B-D)** Doubling time of Jurkat cells, total population growth at the 48 hour endpoint and growth curves are reported. Data shown represents the mean with standard error and each biological replicate shown (n=3). Significance was determined via one-way ANOVA with post hoc Tukey’s multiple comparison test.

### Conditioned media from the lymph node stroma model with flow acts as a chemoattractant for Jurkat egress from the ML model

As the meningeal lymphatic-lymph node pathway is a critical interface for modulation of CNS immune dynamics, we then wanted to directly co-culture Jurkat cells in the ML model and assess egress from the meningeal lymphatics to the lymph node using conditioned media from our lymph node stroma model. We hypothesized that leukemia cells could egress, or exit, the meninges via the meningeal lymphatic vessels, wherein they traffic to the draining lymph nodes and alter immune responses. It is thought that there is bidirectional crosstalk between the brain and the cervical lymph nodes either through the meningeal lymphatic or blood vasculature, thus conditioning lymph nodes with drained fluid from the brain (28). Therefore, activated lymph nodes with increased IFF due to inflammation could stimulate the egress of leukemia cells. To simulate this process in the meninges to lymph node axis, we combined our lymph node stroma and ML models. Our lymph node model consists of a monolayer of LECs beneath a PhotoHA-collagen gel laden with fibroblastic reticular cells (FRCs) as previously described, which are both important stromal components and previously shown to be responsive to IFF (23). Media from our lymph node stroma models under static and flow conditions was then placed underneath the ML model to determine whether the media could act as a chemoattractant and elicit leukemia egress from our ML model (**Figure 3A**). Interestingly, media from the lymph node stroma under flow caused a significant increase in Jurkat egress from the ML model (**Figure 3B**). We also observed that Jurkats remain adhered to the model, but did not cause extensive disruption to the cellular monolayers within the model (**Figure 3C**).

**Figure 3.**
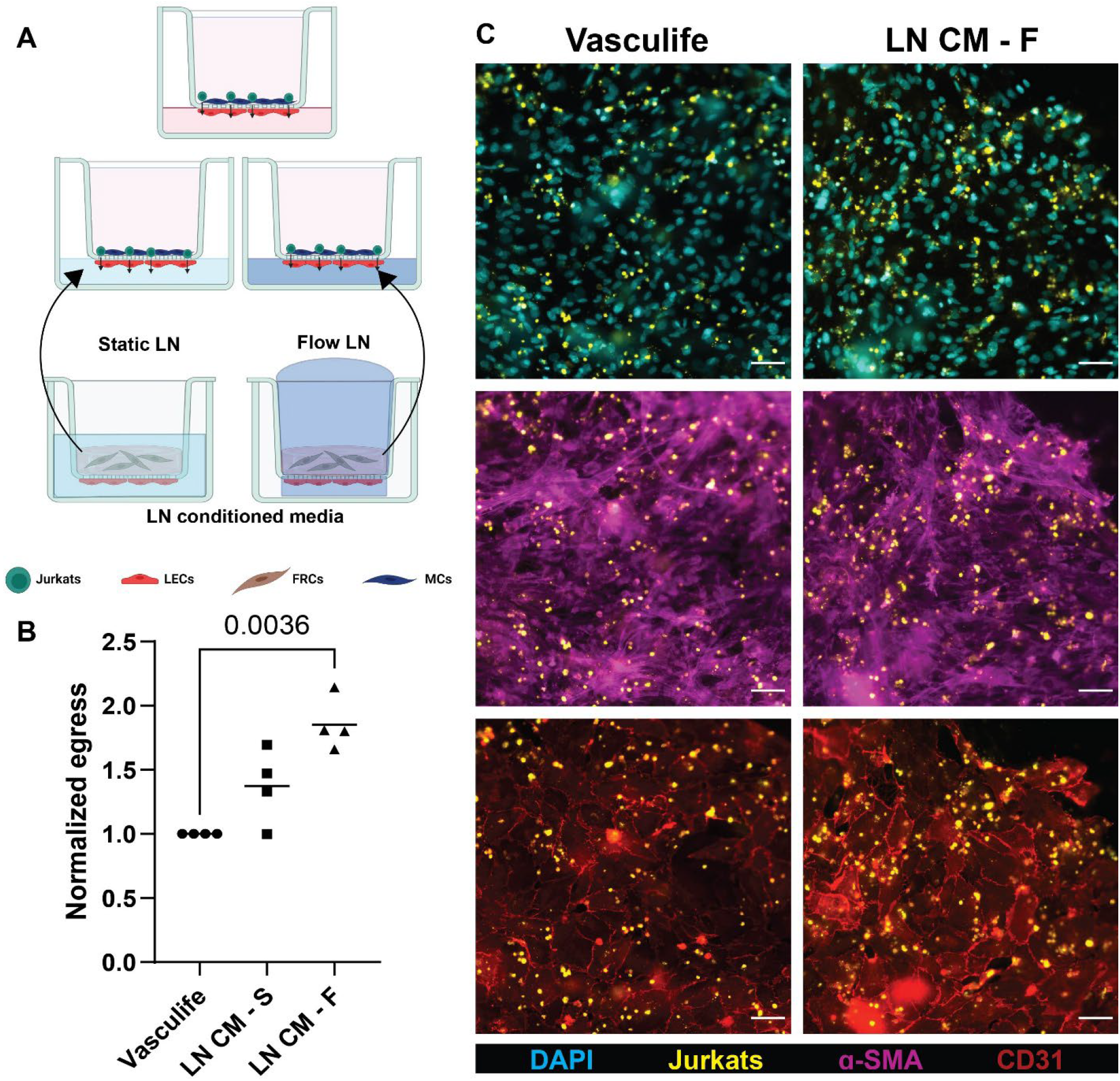
Jurkats egress from the meningeal lymphatic model at higher rates when flow-treated lymph node stroma conditioned media is used as a chemoattractant. **(A)** Lymph node stroma models were cultured under static and flow conditions and the supernatant (denoted as lymph node (LN) conditioned media (CM) with S for static and F for flow) was used as a chemoattractant underneath of the ML model. **(B)** Egress of Jurkat cells from the ML model is reported. **(C)** Representative images show nuclei stained with DAPI (cyan), Jurkat cells labeled with CellTracker Orange (yellow), MCs labeled for alpha smooth muscle actin (magenta), and LECs labeled for CD31 (red). Scale bars are 50 microns. Data shown represents the mean with each biological replicate shown (n=4). Significance was determined via one-way ANOVA with post hoc Tukey’s multiple comparison test.

### Jurkat egress from the lymph node stroma trends upward under flow

Leukemia cells are known to metastasize to the lymph node and replace normal lymphocytes (29). In the context of our results of enhanced egress toward flow-conditioned lymph node stroma, we also wanted to investigate Jurkat behavior within the lymph node microenvironment as metastasis can result in compromised immunity. We previously demonstrated that IFF can enhance inflammatory markers in our lymph node stroma model (25), further impacting leukemia cell behavior. Thus, we explored whether leukemia cells would respond to enhanced fluid flow. As expected, IFF caused a significant increase in Jurkat migration (**Supplemental Figure 1**). Next, we added Jurkat leukemia cells to our lymph node stroma model under static and flow conditions to determine if the presence of flow would impact leukemia migration in the lymph node microenvironment (**Figure 4A**). Although the increase in egress under flow conditions did not reach statistical significance, the trend suggests that leukemia cells may be influenced by fluid flow (**Figure 4B**). Analyzing other metrics, such as leukemia cell proliferation and movement within the lymph node model, may be more impactful. Jurkats were also shown to have enhanced clustering under static conditions (**Figure 4C**), suggesting that co-culture may impact proliferation in addition to migration.

**Figure 4.**
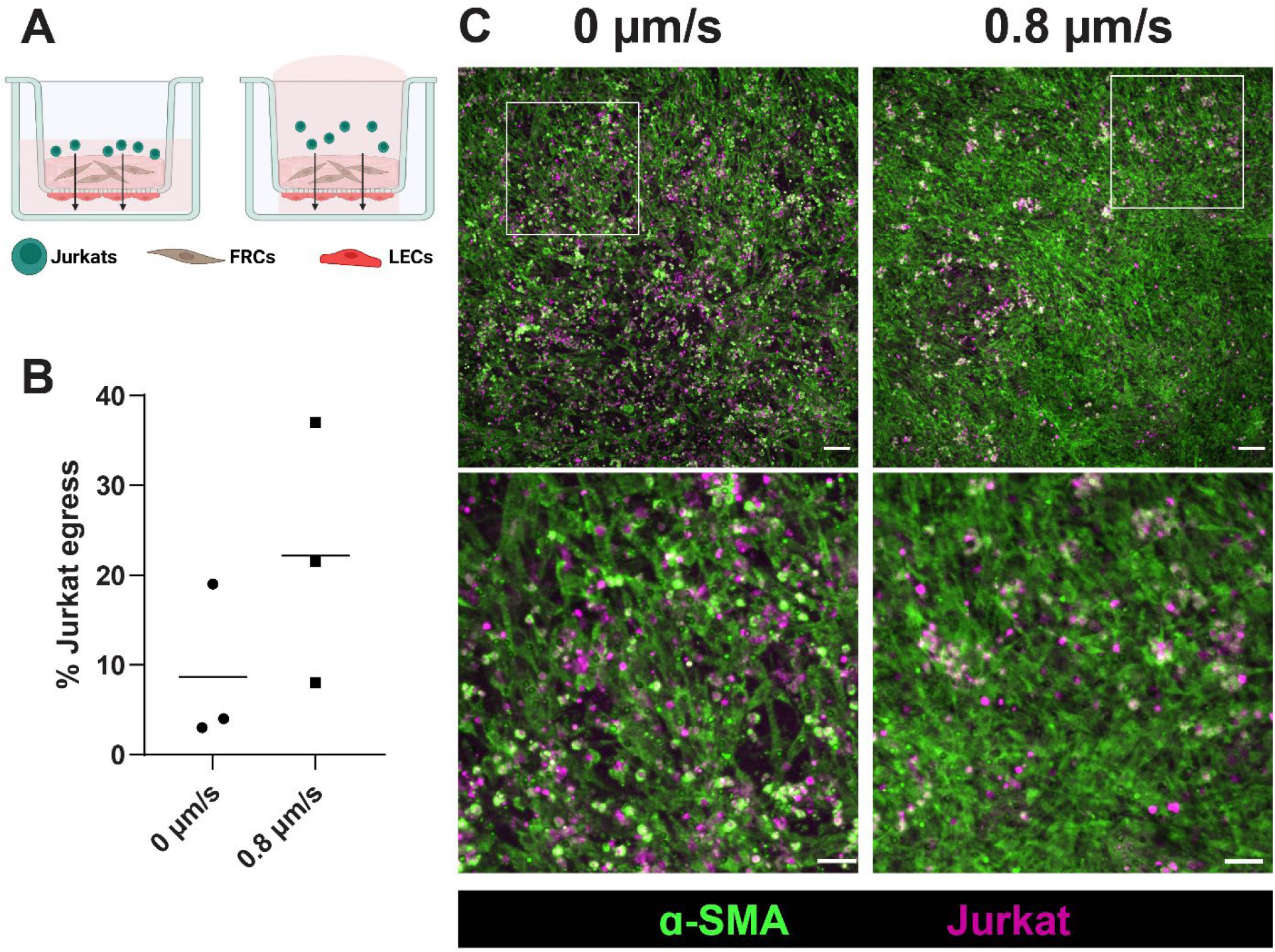
Jurkat egress trends upward in the lymph node stroma under flow. **(A)** Jurkats were added to the lymph node stroma model under static and 0.8 µm/s flow conditions. **(B)** After 24 hours, Jurkats that egressed through the model were located in the lower chamber and counted. **(C)** Representative images show Jurkats labeled with CellTracker Orange (pink) that remained associated with FRCs labeled with alpha smooth muscle actin (green) in the model. Scale bars are 100 microns in the top row and 50 microns in the zoomed images. Data shown represents the mean with each biological replicate shown (n=3). Significance was determined via one-way ANOVA with post hoc Tukey’s multiple comparison test.

**Figure 5.**
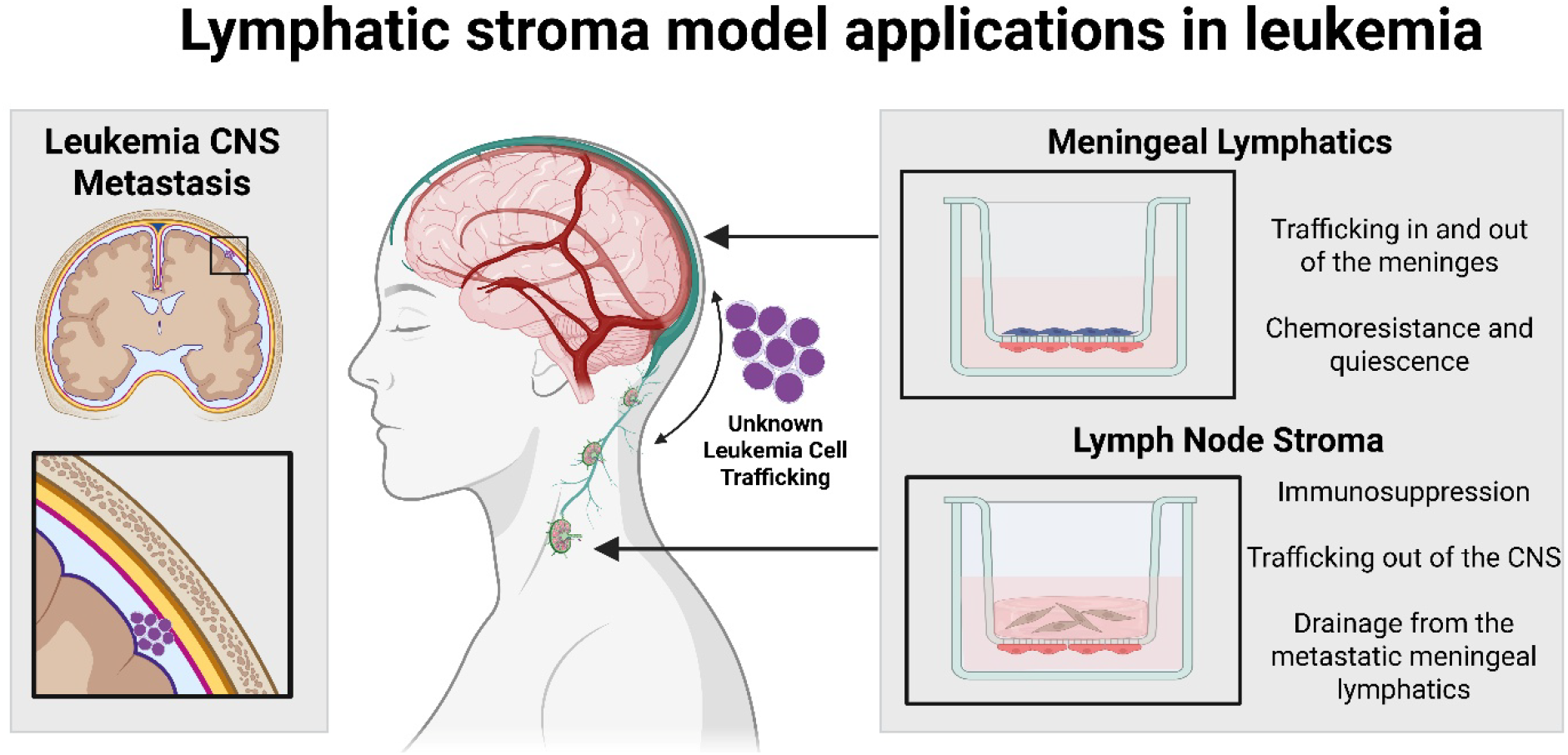
Lymphatic stroma model applications in leukemia. The meningeal lymphatics and lymph node stroma models are useful toward understanding leukemia metastasis, growth, and chemoresistance.

## Discussion

In this study, we have demonstrated that leukemia cell behavior is altered by crosstalk with engineered lymphatic stroma models with a particular focus on the brain to cervical lymph node pathway. Within the last decade, the importance of the meningeal lymphatics in CNS function and cancer progression has gained strong attention, highlighting the need to better understand their function and interactions (12,13). However, the role of meningeal lymphatics in CNS relapse of leukemia, particularly as a potential route for leukemic cell entry into and exit from the CNS, remains poorly understood. Current studies of the brain to cervical lymph node pathway rely on in vivo models and often overlook the meningeal lymphatics, but interest in microfluidic devices to model this pathway has steadily grown (30–32). Thus, developing in vitro models that incorporate the key lymphatics of this pathway and are easily interfaced with other tissues will improve our ability to study leukemia and other diseases with CNS involvement. Within, we investigated the impact of standard of care therapeutics and leukemia cell behavior and response within models representing the meningeal lymphatic and lymph node microenvironment.

Overall, we demonstrated that methotrexate, but not cytarabine, was significantly deleterious to LECs in the ML model. Cytarabine treatment caused a downward trend of LEC coverage, but this was not statistically significant. Methotrexate consistently disrupted LECs and not MCs within the ML model, similar to what previously demonstrated in the presence of taxane chemotherapeutics (10). Generally, methotrexate delivery to the lymphatics is much more efficient when encapsulated in nanoparticles (33,34). However, our ML model directly delivers methotrexate to the cells. In intrathecal therapy, which is a standard component of ALL therapy, methotrexate is injected directly into the cerebrospinal fluid and thus delivered to the meninges. The meningeal lymphatics are actively draining this system and thus, may receive higher doses of methotrexate compared to other lymphatic systems in the body. A disrupted meningeal lymphatic barrier due to methotrexate treatment could result in decreased drainage to the cervical lymph nodes, and thus, an impaired immune response. Alternatively, this disrupted barrier could lead to leukemia cell retention in the meninges, as one route of drainage and trafficking would be disrupted. Thus, the damaged lymphatic barrier could aid in the development of a leukemia cell reservoir in the meninges.

Interestingly, neither chemotherapeutic was disruptive to the MCs. Previous work has demonstrated that MCs enhance leukemia chemoresistance to both methotrexate and cytarabine (8). MCs could secrete various chemokines in response to methotrexate and cytarabine to alter leukemia survival. Thus, exploring the contrast between LEC and MC response to chemotherapy could be an interesting future direction toward understanding this CNS metastatic niche.

We further sought to understand how conditioned media from treated ML models would result in altered Jurkat growth. Interestingly, we saw that conditioned media from the ML model caused the highest trends of Jurkat growth. At 48 hours, Jurkat growth was significantly increased by ML model conditioned media compared to RPMI control. This is particularly notable as the control consisted of fresh media with a higher serum content, pointing to soluble factors secreted by LECs or MCs in the model as responsible for the enhanced growth.

Exploring the pathway of drainage from the CNS to deep cervical lymph nodes, we then examined whether Jurkat cells could egress through the ML model. As the meningeal lymphatics are the primary route for CNS drainage from the meninges, it is plausible that leukemia cells could exploit this route as well to exit meninges. We demonstrated that Jurkat cells were able to egress from the ML model under all conditions. Interestingly, conditioned media from the lymph node stroma model under 0.8 µm/s flow conditions caused enhanced egress. Previously, we demonstrated that 0.8 µm/s flow did not cause changes to the lymph node stroma model morphology or inflammatory chemokine secretion (23). Therefore, alternative soluble factors must be responsible for enhanced egress. LECs and FRCs secrete a number of chemo-attractants that typically impact immune cell migration, egress, and survival under homeostatic conditions (14,35,36). We also demonstrated that a portion of Jurkat cells remain adhered to the ML model. As adherence to MCs is known to enhance leukemia chemoresistance (8), chemoresistance after adherence to the ML model is of great interest for future studies.

Finally, to gain insights on leukemia cell behavior within the lymph node stroma, we incorporated leukemia cells with IFF representative of the microenvironment. Although we showed that Jurkats are flow-responsive when encapsulated in hydrogels alone, once Jurkats were added to the lymph node model, egress trended upward under flow but was not significant. Interestingly, we note that the representative images show clusters of Jurkat cells in static conditions. In cell culture, leukemia cells are non-adherent cells, remain in suspension, and preferentially grow in clusters (37). Therefore, this aggregated state suggests that the static lymph node microenvironment may promote proliferation rather than migration. Further, leukemia cells gradually replace functional immune cells in the lymph node over time (38,39), highlighting the lymph node microenvironment’s capability to support leukemia cell survival. However, this phenomenon has not been extensively studied. To support this, Jurkat leukemia cells have commonly been used as a human T cell substitute in studying the lymph node microenvironment (40–43). Therefore, the behavior of Jurkat cells in non-cancerous contexts can help inform future experiments.

Overall, we have analyzed leukemia cell interactions with the lymphatic system within distinct regions along the brain-to-cervical lymph node pathway. Notably, soluble factors within conditioned media from both the ML model and the lymph node model were able to modulate leukemia cell behavior, specifically in the context of growth and migration, respectively. Our findings underscore the critical need to explore leukemia-lymphatic crosstalk in CNS leukemia and the suitability of our models to help answer key biological questions. Adding additional complexity to the models is of great interest in future work. For instance, the ML model could be cultured in artificial CSF or interfaced with a brain parenchyma model. On the other hand, the lymph node model could incorporate lymphocytes prior to metastasis to examine anti-tumor immunity. Both the lymph node and ML models could be interfaced into a device to form a complete meninge-to-lymph node pathway model. Additionally, these models could be applied toward other solid and hematological malignancies that are known to metastasize to the leptomeninges, including breast cancer, lung cancer, and lymphoma (44). With easily adaptable models using commercially available components, our hope is to establish these models as widely applicable and modifiable tools for use in understanding CNS leukemia and other pathologies of interest.

## Conclusion

Herein, we have demonstrated versatile applications of engineered lymphatic stroma models in advancing our understanding of CNS leukemia relapse and subsequent egress to the lymph node. We highlight the meningeal lymphatics as an important consideration during CNS-directed chemotherapy for leukemia. Additionally, we showed that enhanced interstitial fluid flow within the lymph node microenvironment facilitated leukemia egress from the meningeal lymphatics. Future work will focus on elucidating which soluble factors are involved in the crosstalk between leukemia cells and CNS lymphatic systems to enhance our understanding and treatment of CNS leukemia.

## Data Availability

The data underlying this article will be shared upon reasonable request to the corresponding author.

## Acknowledgements

JH was supported by the Institute for Critical Technology and Applied Science (ICTAS) at Virginia Tech. Schematics (Figure 1A, 2A, 3A, 4A, 5) were created with Biorender.

