## Supplemental Figure 1 for "Engineered lymphatic stroma model applications in central nervous system leukemia"

### Supplemental Material

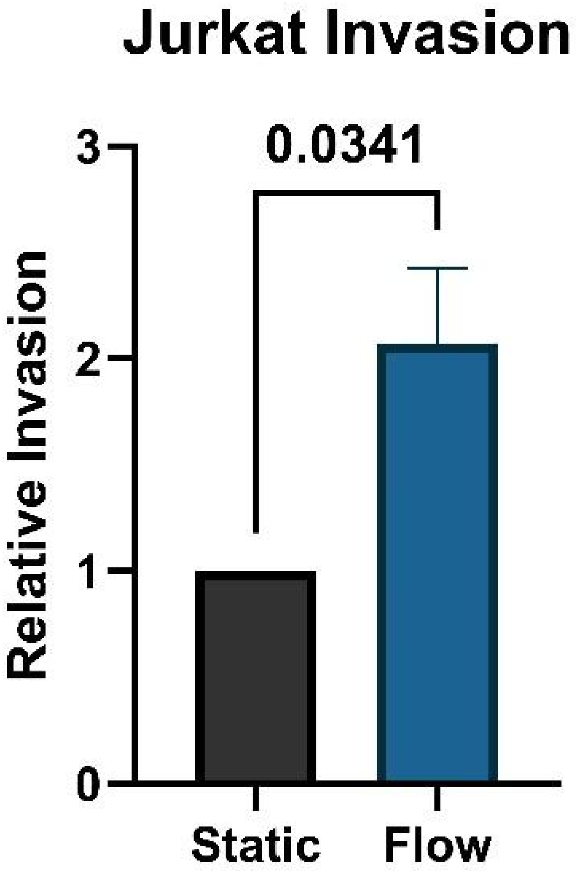


**Supplemental Figure 1**. Jurkats invade into tissue culture membranes at significantly higher rates under interstitial fluid flow (n=3). Statistical significance was determined by students' unpaired T-test with significance set to $p<0.05$.
